# Functionally distinct dopamine domains in the hippocampus

**DOI:** 10.1101/2024.01.31.578232

**Authors:** Muneshwar Mehra, Gerardo Molina, Md Tarikul Islam, Zahra M. Dhanerawala, Eleonora Bano, Edward B. Han

## Abstract

Numerous studies have identified dopamine signaling in the hippocampus as necessary for certain types of learning and memory. Since dopamine in the striatum is strongly tied to rewards, dopamine in the hippocampus is thought to reinforce reward learning. Despite the critical influence of dopamine on hippocampal function, little is known about dopamine release in the hippocampus or the specific ways dopamine can influence hippocampal function. Based on the functional complexity of hippocampal circuitry, we hypothesized the existence of multiple dopamine signaling domains. Using optical dopamine sensors, two-photon imaging, and head-fixed behaviors, we identified two functionally and spatially distinct dopamine domains in the hippocampus. The “superficial” domain (cell somata and apical dendrites) showed reward-related dopamine transients early in Pavlovian conditioning but were replaced by “deep” domain transients (basal dendritic layer) with experience. These two domains also play distinct roles in a hippocampal-dependent, goal-directed virtual reality task where mice use exploratory licks to discover the location of a hidden reward zone. Here, positive dopamine ramps appeared in the superficial domain as mice approached the reward zone, similar to those seen in the striatum. At the same time, the deep domain showed strong reward-related transients. These results reveal small-scale, anatomically segregated, dopamine domains in the hippocampus. Furthermore dopamine domain activity had temporal-specificity for different phases of behavior. Finally, the subcellular scale of dopamine domains suggests specialized postsynaptic pathways for processing and integrating functionally distinct dopaminergic influences.

## Introduction

Dopamine has been intensively studied in the striatum where it is broadly associated with both rewards and movement^1–5^. Functional differences in dopamine signals coarsely align with anatomy; movement correlated dopamine is more prevalent in dorsal striatum which is primarily innervated by afferents from substantial nigra pars compacta (SNc), while reward prediction error (RPE) and motivation are more prominent in the ventral striatum which is primarily innervated by the ventral tegmental area (VTA)^6^.

Far less is understood about dopamine’s roles outside of the striatum, where lower dopamine levels make accurate measurement difficult. Numerous studies have found that dopamine is necessary for hippocampal-dependent learning, perhaps by stabilizing long-term plasticity or memory consolidation^7–9^. However, the exact role dopamine plays is unclear since crucial information is missing: when and where is dopamine released in the hippocampus?

Here we address two questions. First, what is the functional role of dopamine in the hippocampus? Does it represent reward or RPE similar to dopamine the ventral striatum? Another distinct possibility is that dopamine conveys movement-related signals as in the dorsal striatum. Movement is critical to hippocampal function as it defines network state, with periods of locomotion marked by place cell firing, while immobility switches the network to replay and consolidation. Furthermore, growing evidence identifies specialized neuronal circuits specifically active during periods of movement and immobility^10,11^.

Second, what is the spatial scale of dopamine domains? Dopamine is a volume transmitter, with no pre- to postsynaptic alignment; however, the size of functional domains could vary by orders of magnitude, from single microns to millimeters^12^. Defining the spatial scale of dopamine domains is critical for interpreting the results of fiber photometry experiments, which have no optical sectioning ability but instead take the mean of fluorescence over hundreds of microns.

## Results

To precisely measure dopamine dynamics with high spatio-temporal resolution, we recorded with optical sensors, GRAB-DA1h, 2h, and 2m^13^ using two-photon imaging to image a volume of tissue with multiple axial planes in head-fixed mice during behavior (Fig. 1A, B). In a Pavlovian task with randomly timed rewards, we paired a conditioned stimulus (CS, sound cue) with water reward (UCS, Fig. 1C). Early in training, while recording with high affinity sensors GRAB-DA1h and 2h, we found layer-specific dopamine responses, with small increases in the superficial domain (corresponding to the apical dendrite and pyramidal cell layers) (Fig. 1D). It was important to align mice by stage of learning to consistently observe reward-related transients in the superficial domain, due to variability in when mice learned the association (Fig. 1E). The magnitude of these transients peaked around the time that animals learned the association between CS and UCS and then rapidly declined; however, even with this alignment, the small magnitude of these transients remained problematic. We also tested the lower affinity sensor (GRAB-DA2m) and surprisingly found larger superficial transients in early training. To quantify reward-related transients, we combined results from GRAB-DA1h, 2h, and 2m, and aligned in CS-triggered averages (Fig. 1F,G, S1A). We found a significant increase in CS-triggered superficial dopamine with no change in deep dopamine (corresponding to the basal dendritic layer). Dopamine levels in controls (dopamine insensitive version of the reporters, GRAB-DAmut, or EGFP) did not change (Fig. 1H). Thus dopamine in the hippocampus is functionally organized in restricted spatial domains defined by anatomy, with reward-related transients in the superficial domain during early Pavlovian conditioning.

**Figure 1.**
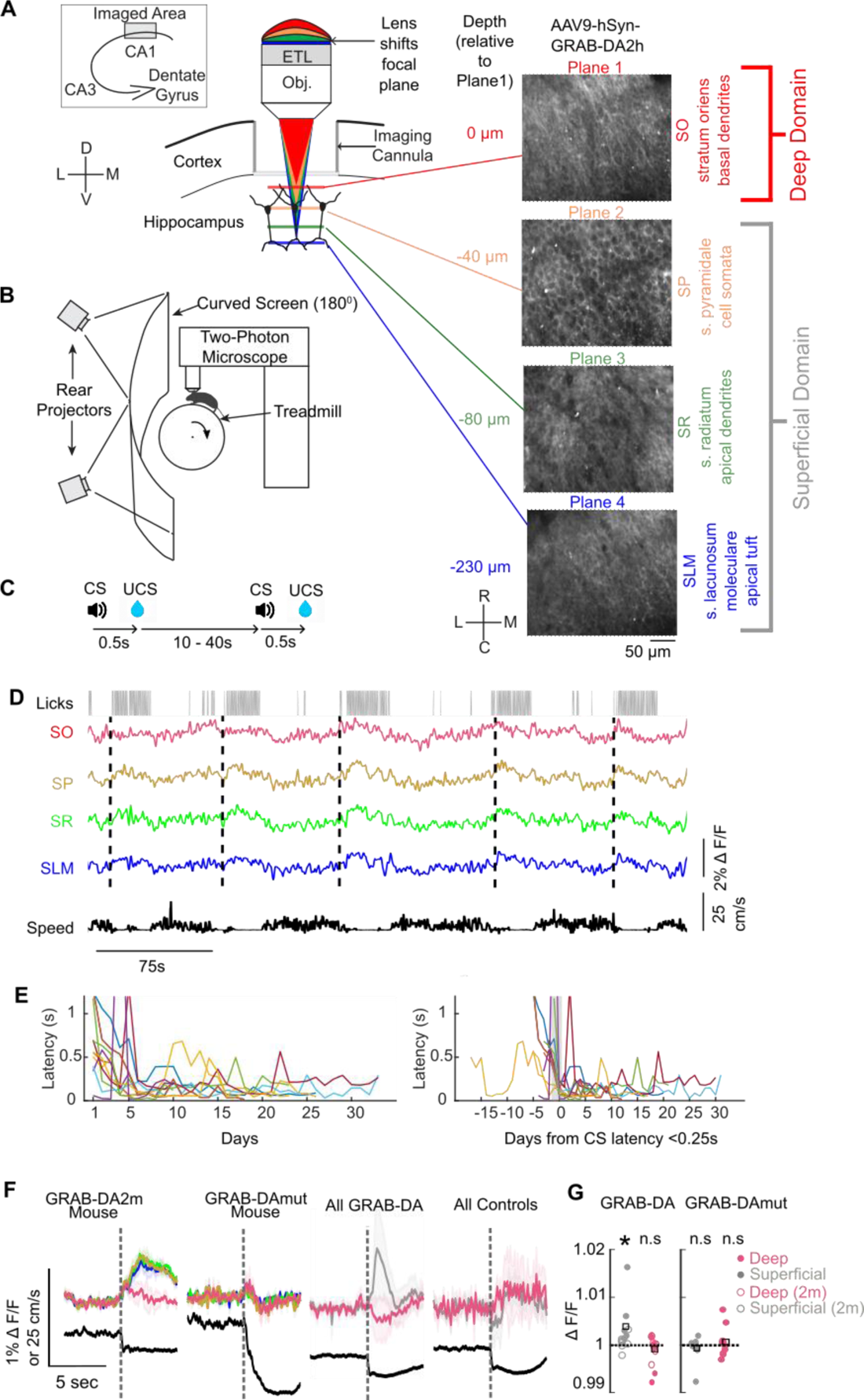
Reward-related dopamine in superficial domain during early Pavlovian conditioning. **A,** Schematic of imaging. Inset, approximate imaged area in dorsal CA1. Electric tunable lens (ETL) allows sequential imaging at different focal planes corresponding to anatomical layers. In some subsequent panels, results from SP, SR, and SLM are averaged together as “Superficial Layers” or “Superficial”. **B,** Schematic of virtual reality (VR) setup. Projection screen not used for random reward experiments. Not shown, lick spigot for water rewards. **C,** top, schematic of reward delivery at random intervals. bottom, raw data from well-trained mouse showing fluorescence in different cell layers, along with CS, licks and speed of treadmill. **D,** CS-triggered average of fluorescence and speed over four imaging sessions in example GRAB-DA2h expressing mouse (left) and control GRAB-DAmut (right). Dotted line = CS. **E,** Left, Lick latency after CS of individual animals over days. Right, lick latency aligned by first day when lick latency after CS < 0.25s. **F,** CS-triggered dopamine signal by layer in example GRAB-2h mouse and control GRAB-DAmut. Locomotion in black at bottom. **G**, Superficial layer dopamine was similar so was averaged into the “superficial” domain. CS-triggered dopamine signal by domain for all experimental and control mice. **H,** CS-triggered dopamine signal was significantly greater in superficial domain and unchanged in deep domain and controls.

It remains unclear why the low affinity sensor shows higher superficial domain signal. One possibility is saturation of the high affinity sensor, although this seems unlikely based on the low concentration of dopamine in the hippocampus^14^ in comparison to the striatum, where these sensors are most often used.

Surprisingly, these reward-related superficial domain transients did not persist throughout continued behavior. Shortly after the amplitude of superficial transients peaked and mice learned the CS-UCS association, superficial transients began to decrease and deep domain transients increased in magnitude (Fig. 2A, B, S2A). Focusing on CS-triggered dopamine in well-trained mice, we found layer-specific dopamine responses (Fig. 2C-E, S2B). In the deep domains, positive dopamine transients coincided with CS presentation and reward. In contrast, dopamine in the superficial domain did not change (Fig. 2F), nor did dopamine levels in controls. Thus dopamine domains signal reward-related activity at different phases in Pavlovian conditioning, with the superficial domain transients early in training and deep domain transients late in training.

**Figure 2.**
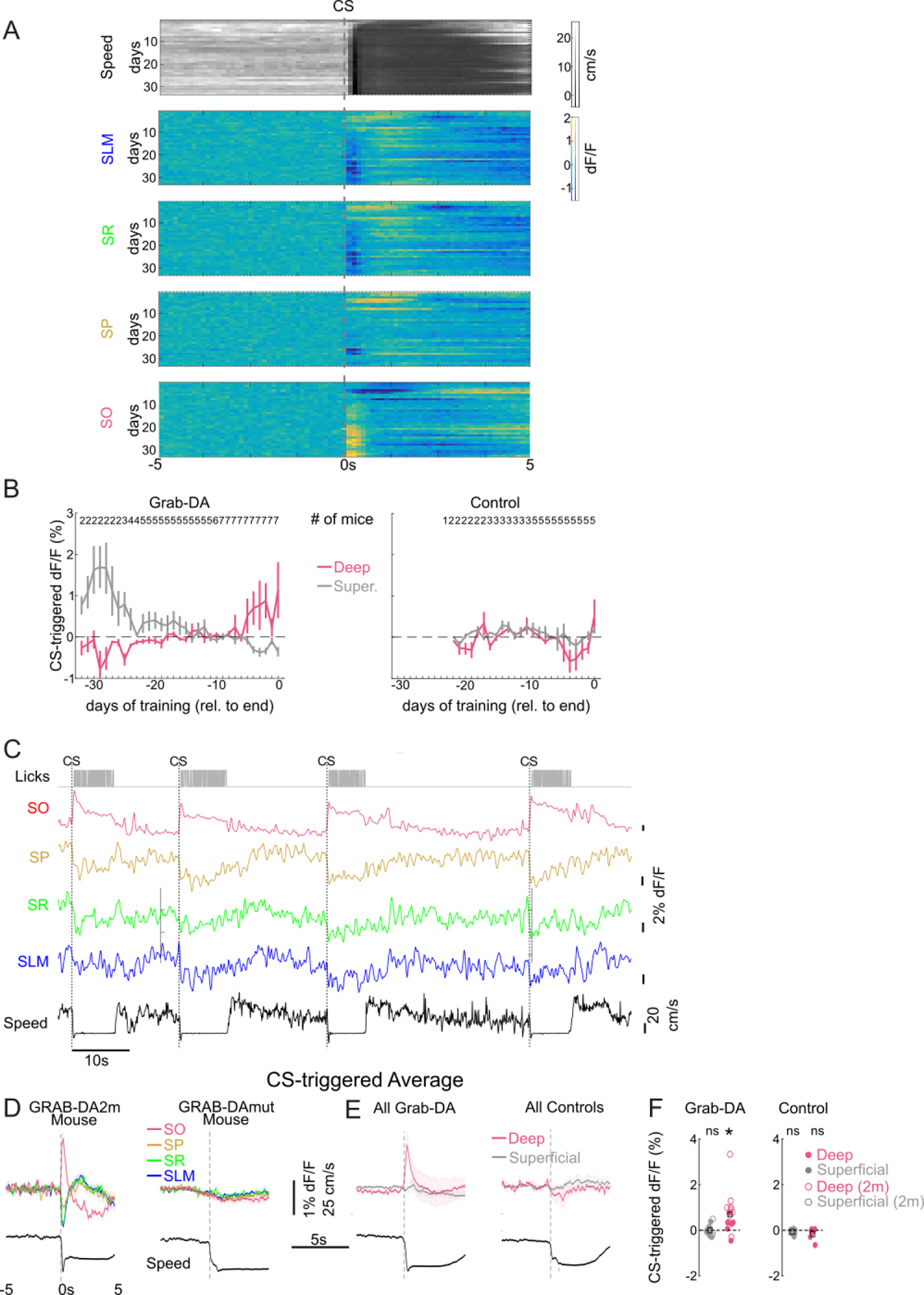
Reward-related dopamine transients shift from superficial to deep domain over experience. **A,** Speed and dF/F dynamics over days of training in an example mouse. **B,** CS-triggered dopamine transients in deep and superficial domains over experience in all experimental mice (left) and controls (right). Numbers at top indicate number of mice for each day. **C,** Late training, raw data from well-trained mouse showing fluorescence in different cell layers, along with CS, licks and speed of treadmill. **D**, CS-triggered dopamine signal by layer in example GRAB-2h mouse and control GRAB-DAmut. **E**, CS-triggered dopamine signal by domain for all experimental and control mice. **F,** CS-triggered dopamine signal was significantly greater in deep domain in late training and unchanged in superficial domain and controls.

Reward-related dopamine transients can convey RPE signals in the striatum and our result showing CS-triggered dopamine transients in the hippocampus is, so far, consistent with this role^2^. To directly test if deep domain dopamine transients in well-trained mice had RPE properties, we inserted a small number of reward omission trials (Fig. 3A), where dopamine levels should decrease based on negative prediction error. However, we found no change in the amplitude of deep dopamine transients with reward omission, nor did dopamine levels change in the superficial domain (Fig. 3B-E, S3A). At ∼1s after CS, the amplitude of deep dopamine decreased in omission trials, however this decreased dopamine is likely driven by locomotion changes since mice only briefly stop after reward omission, as opposed to longer stopped licking with rewards. We also tested the effect of reward addition by doubling rewards, either by the addition of a second, uncued reward after the cued reward, or signaled by a double CS. Doubling the expected reward did not change dopamine levels (S3B). Thus, we found insensitivity to unexpected changes in rewards in deep dopamine transients in well-trained mice, indicating that this distinct dopamine signal does not represent RPE. We did not test early in training so it remains to be seen if superficial domain dopamine has any reward prediction properties.

**Figure 3.**
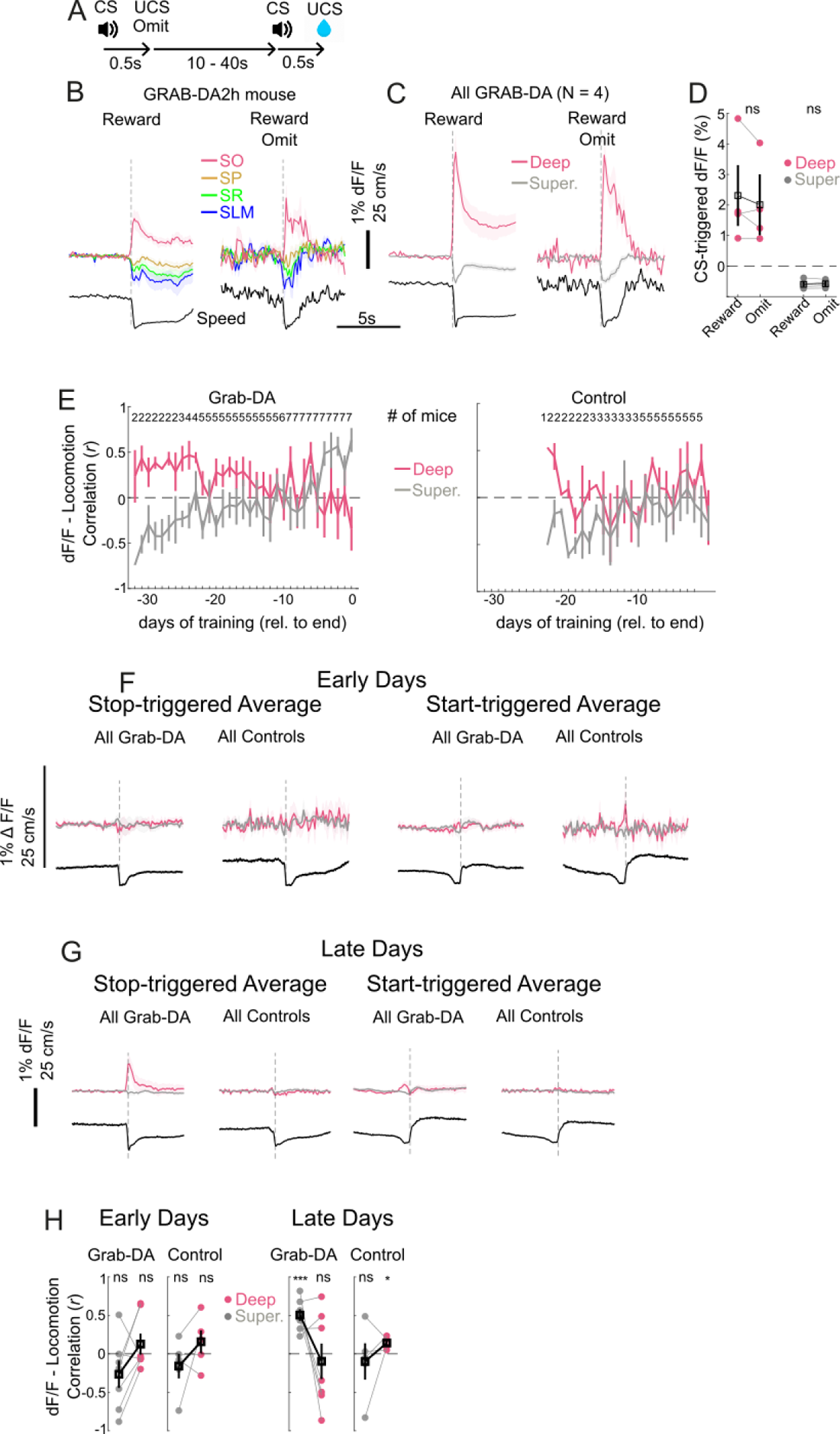
No effect of reward omission on deep dopamine and locomotion correlation of superficial dopamine. **A,** Schematic of reward omission. **B,** CS-triggered average of fluorescence for rewarded (left) and unrewarded CS (right) from same example mouse. Note that stopped period during reward omission is shorter than rewarded with a corresponding decrease in the late phase of deep domain fluorescence. **C,** CS-triggered average for reward and reward omit from all experimental mice. **D,** No difference in CS-triggered dopamine transients for reward and reward omit. **E,** Correlation and anti-correlation between dopamine and locomotion over experience. Numbers at top indicate number of mice for each day. **F**, Early training locomotion start-triggered and stop-triggered average of dopamine in experimental and control mice. **G**, Late training locomotion start-triggered and stop-triggered average of dopamine in experimental and control mice. Note the visual correlation between superficial dopamine. **H**, Dopamine signal in superficial domain was significantly correlated with locomotion during late training.

We have described reward-related dopamine activity but locomotion also co-varies with rewards. During behavior, mice run in between reward presentations, typically starting a few seconds after reward and continuing until the CS, when they stopped and licked (Fig. 2A). Thus we investigated the relationship between locomotion and dopamine activity. Over the time course of Pavlovian conditioning, superficial domain dopamine was anticorrelated to locomotion early, and then became more correlated over time (Fig. 3E, S3C). The early anticorrelation with locomotion was consistent with reward-related transients that coincided with stopping to lick rewards. In contrast, deep domain dopamine followed the opposite trend, going from correlation to anticorrelation. To examine the relationship between locomotion and dopamine, we made start and stop-triggered averages of dopamine (Fig. 3F,G), since these behaviors seemed to drive overall correlations. Stop-triggered averages look very similar to reward-triggered averages because most of the stops are due to reward, with relatively few spontaneous stops. In late days of training, the superficial dopamine domain showed start-triggered averages that qualitatively correlated with the locomotion start signal, while superficial dopamine also decreased at stops (Fig. 3F). Indeed the superficial dopamine domain signal showed significant correlation with locomotion speed, when calculated over the entire session, late in training (Fig. 3H), indicating that this domain of dopamine can represent reward early in training and switch to a movement related signal later in the same behavior.

To study dopamine dynamics during more complex, goal-directed learning, we developed an instrumental learning paradigm, the VR hidden reward zone task (HRZ, Fig. 3A,B). Previous work found characteristic ramping up of dopamine in the striatum prior to reward, when the reward occurs at a predictable time or place^1–3,6,15^. The function of this ramp in the striatum remains controversial, either signaling increasing value/motivation^1,15^ or acting as RPE^2^. Regardless of its function, we asked whether this learning-related signature of striatal dopamine is also present to the hippocampus, and if so, is it also targeted in a domain-specific manner.

In the HRZ task, mice run on a treadmill to control their position on a virtual track. On reaching the end of the track, they experience a random duration dark time (2 - 4s), and then begin another trial at the beginning of the track. Mice are trained to lick the water reward spigot as an exploratory behavior. A lick when the mouse is in the unmarked reward zone results in a CS and subsequent water reward. After discovering the location of the reward zone, mice refine their lick pattern closer towards the reward zone. At the end of each epoch, there are three unrewarded probe trials, and the reward zone moves to another hidden location.

Indeed, we found hippocampal dopamine ramps that were domain-specific. Surprisingly, we identified both positive and negative ramps during reward approach. In the superficial domain, there was a positive ramp prior to CS, while the deep domain had a negative ramp (Fig. 4B-E, S3A). The superficial domain positive ramp was qualitatively similar to those seen in ventral striatum, starting when mice began navigating the track and stopping shortly before entering the reward zone. However, one might expect the deep domain to have positive ramps since it had reward-related transients in HRZ, in analogy to ventral striatal dopamine, which has positive ramps and is strongly tied to rewards^2,15^. We note that negative dopamine ramps or negative firing ramps in dopaminergic neurons have been seen by multiple groups, although their function remains unclear^3,15^. Dopamine ramps were well fit by linear regression in experimental animals but not in controls (S3 B,C). There was no ramping of dopamine prior to CS in the Pavlovian conditioning task (S3 D), presumably because the variable inter-reward intervals prevent predictable reward expectation.

**Figure 4.**
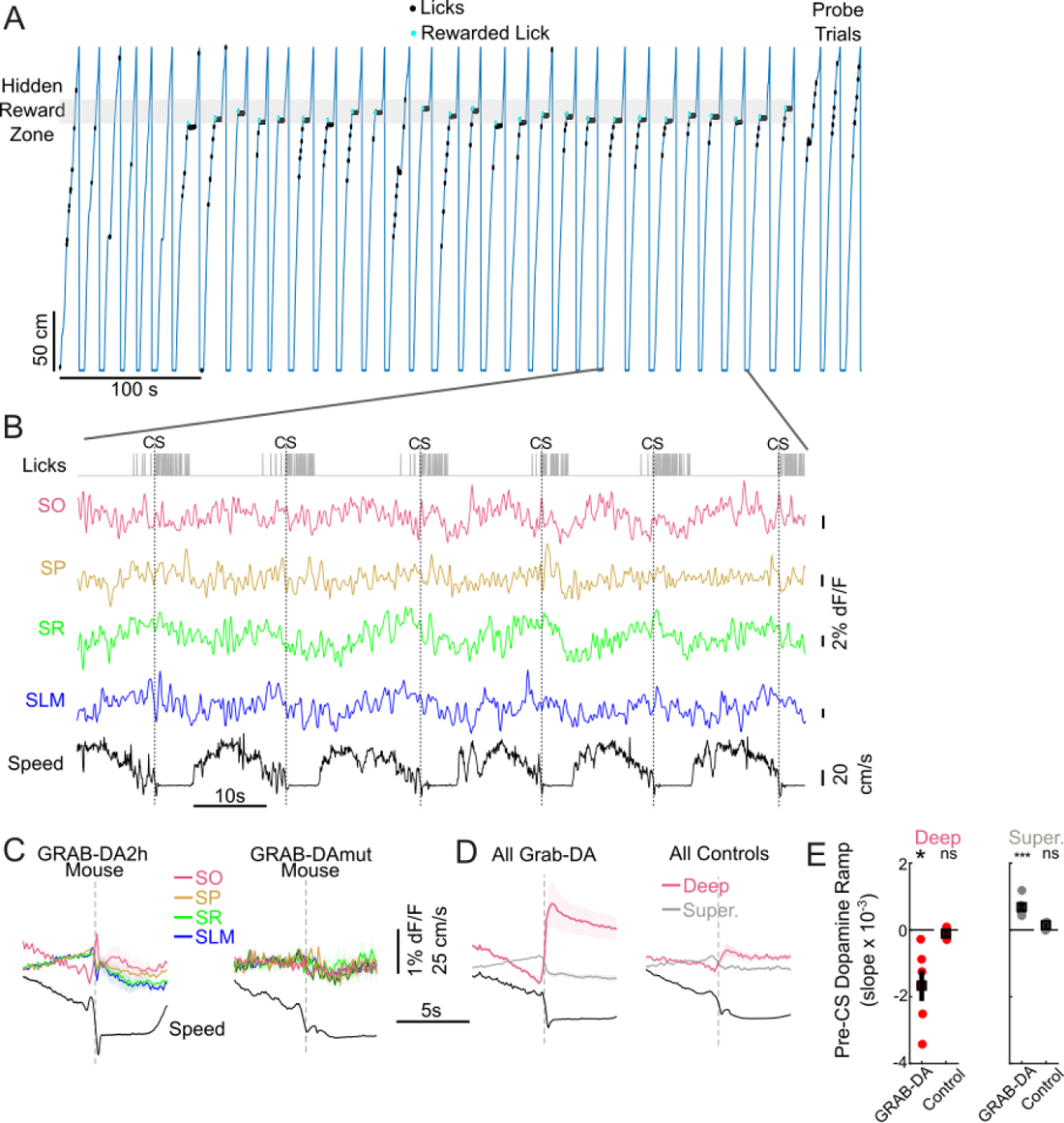
Dopamine ramps in a VR hidden reward zone task (HRZ). **A,** HRZ behavior. Licks in the uncued, hidden reward zone trigger CS and water reward. Trials are separated by random duration dark time. Blue line is mouse’s position on virtual track. At the end of each epoch, there are three unrewarded, uncued probe trials. **B,** Licks, plane fluorescence, and speed expanded from marked period in **A**. **C,** CS-triggered average from example experimental and control mouse. **D,** CS-triggered average from all experimental and control mice. **E,** Superficial domain showed positive dopamine ramps while deep domain had ramping during reward approach, based on slopes of linear fit.

The positive dopamine ramp in the superficial domain in the HRZ task was also unexpected because in Pavlovian conditioning, this dopamine signal is locomotion-correlated. However, here the positive dopamine ramp is anticorrelated with locomotion, specifically prior to reward, as mice slow down while approaching the reward zone (Fig. 4C,D, S4A,E). To further define the relationship between locomotion and superficial layer dopamine in the HRZ task, we looked at start-triggered averages and whole time series correlation between dopamine and locomotion. Similar to late Pavlovian conditioning, we found that superficial layer dopamine increased at locomotion start and was significantly correlated with locomotion over the entire session (S4 G). Thus dopamine levels in the superficial domain during HRZ correlates well with locomotion, except during approach to expected rewards.

## Discussion

Our results reveal anatomically-defined, functional dopamine domains in the hippocampus. During Pavlovian conditioning, there were reward-related transients in the superficial domain (corresponding to the apical dendrite and cell bodies) early in training. Around the time mice learned the association between CS and reward, these superficial domain transients shrank and were replaced by deep domain transients (corresponding to the basal dendritic layer). We identify a restricted spatial scale for these domains, as small as ∼100μm (corresponding to the thickness of the SO layer), with functionally and spatially distinct domains in adjacent layers. Importantly domains at such spatial scales cannot be observed using fiber photometry or miniscopes. Since these domains are axially segregated, single photon approaches with no optical sectioning ability will not be able to resolve them. Our findings raise the possibility of identifying other small dopamine domains in other brain regions when measured at sufficient resolution during behavior.

The finding of distinct dopamine signals in close apposition was striking and naturally raises the question of their anatomical and functional origins. Both dopaminergic midbrain (VTA and SNc), and LC dopaminergic neurons project to the hippocampus. Studies of layer-specific targeting by VTA dopaminergic axons are contradictory with one finding primarily SO targeting^16^, and another finding preferential SP targeting^17^, although both found axons in all layers. LC axons (as subset of which are dopaminergic) also seem to target all layers^18,19^. Calcium imaging of axonal inputs provide complementary results to our direct imaging of dopamine in the hippocampus. VTA dopaminergic inputs to the hippocampus were found to have ramping activity prior to reward during track-running^21^, similar to the signal we saw in the superficial domain in the HRZ task. How inputs from these three areas, VTA, SNc, and LC, are integrated to produce the rich dopamine dynamics we identify remains unclear. Yet another complicating factor is that even within VTA and SNc, there is strong functional heterogeneity along the medio-lateral axis^3,20^. Dopamine domains could also arise through local modulators of axonal dopamine release such as acetylcholine, independent of dopaminergic somatic activity^1,22^. Dopamine waves may also contribute to the development of hippocampal domains. Previous work in the striatum identified waves of activity that propagated in opposite directions for instrumental and Pavlovian tasks and these could play a role in differences in dopamine release between our Pavlovian and HRZ tasks^23^.

The network effect of these dopamine domains require further investigation but our findings show that postsynaptic neurons can see differing levels of dopamine with subcellular resolution, rather than a uniform global signal. These restricted domains imply correspondingly fine scale postsynaptic mechanisms to process distinct reward and locomotion signals with subcellular or microcircuit resolution. One possibility is preferential activation of specific classes of interneurons to exert dopaminergic influence on the circuit, for example, OLM interneurons have dendritic arbors completely within SO. Dopamine has also been proposed to tag specific synapses for consolidation through late long-term potentiation, and here could act by selectively tagging SO synapses activated at rewards^24,25^.

Finally the insensitivity of deep dopamine to reward omission late in Pavlovian conditioning indicates that it does not function as a rapid feedback signal like RPE but is more consistent with an automatic or habitual behavior. This dopaminergic activity is somewhat similar to that seen in a class of macaque dopaminergic SNc neurons that slowly learn reward values and then become insensitive to reward omission^26^. The RPE properties in HRZ and in superficial dopamine in early Pavlovian conditioning remain untested. Further future experiments will elucidate the origins of these complex signaling pathways and their specific roles in hippocampal processing. Regardless of how these domains arise, our results reveal the fine scale of dopamine signaling, functionally specialized pathways, and behavioral dynamics of hippocampal dopamine signaling.

## Methods

Procedures were similar to those previously described^10,27^ Animals All experiments were approved by the Washington University Animal Care and Use Committee. Wild-type mice of both sexes on a C57BI/6J background were used for this present study. Mice were 2-3 months old at the time of surgery.

### Viral Injection and hippocampal window implantation

Mice were anesthetized with isoflurane, after which a 0.3 mm craniotomy was opened above the left cortex. Subsequently, viral vectors, including AAV-hsyn-GRAB_DA1h ( mice 156 and 157), AAV-hsyn-GRAB_DA2h (2.6 × 1013, diluted 1:1-5 with PBS, mice 167, 168, 169, 171), AAV-hsyn-GRAB_DA2m (2.6 × 1013, diluted 1:1-5 with PBS, mice 220, 221, 222), AAV-hsyn-GRAB_DA-mut (2.0 × 1013, diluted 1:1-5, mice 170, 179 and 181), AAV2-CAG-Flex-eGFP-WPRE-bGH 8.2 × 1012, diluted 1:10, mice 158) were pressure injected (∼50nL) through a beveled micro-pipette targeting dorsal CA1 (−2.4 mm posterior, −1.6 mm lateral, −1.3 mm ventral, relative to Bregma). Mice were water scheduled 2-3 weeks post virus injection. Afterward, an imaging cannula (2.8 mm diameter) was implanted above CA1 by aspirating the overlying cortex. Mice were allowed to recover at least two weeks after surgery before the commencement of behavioral training and imaging.

### Imaging

Imaging was conducted using a laser scanning two-photon microscope (Neurolabware) equipped with an electrically tunable lens (ETL; Optotune, EL-10–30-NIR-LD), enabling rapid focal plane shifts. The microscope control and data acquisition were conducted using Scanbox (Neurolabware). The microscope was specifically configured to capture dopamine activity within the dorsal CA1 region of the hippocampus. The z-axis was scanned to obtain 3-4 axial planes, corresponding to different layers: Stratum Oriens (SO), Stratum Pyramidale (SP), Stratum Radiatum (SR), and Stratum Lacunosum Moleculare (SLM). Identification of specific strata was achieved through distinct features. SP was recognized by the presence of cell bodies, while SLM was identified based on depth and dark areas corresponding to blood vessels. Imaging depths for different strata were as follows: SO at −50 to −80 μm, SP at −110 to −130 μm, SR at −170 to −200 μm, and SLM at −300 to −350 μm. The field of view (FOV) in the x-y plane was approximately 350 × 350 μm. The total frame rate of imaging was 31.25 Hz, resulting in a per-plane sampling rate of 10.4 Hz for a three-plane recording and 7.8 Hz for a four-plane recording.

The GRAB-DA signal was captured at 920nm, with power ranging from 20 to 39 mW per plane after the objective. Higher powers were used for deeper planes to ensure adequate fluorescence intensity. Furthermore, GRAB-DAmut, a variant with dimmer fluorescence, required higher powers in the range of 39 to 51mW to match intensity levels with GRAB-DA mice. Additionally, EGFP (Enhanced Green Fluorescent Protein) was employed as a control and imaged at 9mW.

The experimental setup involved head-fixing mice above a cylindrical Styrofoam treadmill mounted on a 3D-printed base. Rotational velocity was tracked using a rotary optical encoder, and licking behavior was measured through an electrical contact circuit. The conditioned stimulus (CS) was provided by the click of a dummy solenoid located behind the mouse. Reward delivery, administered by a second solenoid, was sound-insulated and positioned several feet away outside the recording rig to ensure mice could not hear the reward solenoid activation.

### Random Reward

In the Pavlovian task, mice underwent habituation by running on a treadmill without water reward delivery for 1-2 weeks. During this period, the animals were in the dark without a virtual reality (VR) world. Following habituation, conditioned stimulus (CS) and unconditioned stimulus (UCS) pairs were presented at random interval inter-trial intervals ranging from 15 to 45 seconds. Additionally, a designated no-reward delay window of 2 minutes was introduced before the initiation and after the conclusion of each session. After the delivery of a water reward, it remained available on the metal tube until consumed at any point during the behavioral session, with most mice quickly associating CS with UCS. Each session, lasting 20-24 minutes, included reward doubling trials (3-4 trials), where a second reward was added 0.5 seconds after the first, either CS-cued or without a cue. As no significant differences were observed, conditions were combined. The experimental phase continued for 2–5 weeks. Omission experiments, conducted in a separate block after random reward training in a subset of animals, involved two randomly inserted reward omissions in each session.

### VR track running behavior

Following completion of the Pavlovian task, mice were transitioned to a virtual environment controlled by ViRMEn (Virtual Reality Matlab Engine, Aronov and Tank, 2014), featuring a visual linear track with both local and distal landmarks, measuring either 180 or 270 cm in length. In this context, mice traversed the track, experiencing a brief dark interval of 2-4 seconds upon reaching the far end before initiating the subsequent trial at the track’s beginning. In the initial stages of training, water rewards were provided when the mouse licked a contact sensor and moved forward a minimum distance of 5 cm. As mice consistently demonstrated licking behavior to receive rewards, a progression occurred wherein rewards were exclusively delivered when the animals occupied a larger uncued reward zone or multiple smaller reward zones. The complexity was then gradually reduced, decreasing the number and/or size of reward zones until a single 10 cm reward zone remained. After achieving success in 20-27 consecutive trials, three probe trials without rewards were introduced, followed by the relocation of the reward zone. These zones were strategically chosen from early, middle, or late sections of the track, ensuring that sequential reward zones did not originate from the same section. The delineation of early, middle, and late sections was based on specific track intervals (67-86, 101-120, 135-154 cm for the 180 cm track), and the center of the reward zone was randomly selected within these intervals. The presented data reflect observations from well-trained animals capable of consistently completing 2-4 epochs per imaging session lasting 20-24 minutes.

### Data Analysis

Data were analyzed using custom programs written in Matlab (MathWorks). Initial to the analysis, the images underwent motion correction through the application of Suite2p software package (Pachitariu et al., 2016), employing cross-correlation registration and rigid translation of individual frames. To identify Regions of Interest (ROIs) for fluorescence measurement, we employed the mean projection of the recording to create a distinct outline of the landmarks. The ROIs were manually defined using the Matlab ‘roipoly’ function, and fluorescence was subsequently extracted by calculating the average intensity across time for all pixels within each ROI. Ensuring consistency across days, the same approximate region of interest was imaged using local landmarks such as blood vessels. Additionally, a gaussian window with a duration of 0.5 seconds was applied to smooth the fluorescence data.

Locomotion was measured with a rotary optical encoder (Yumo E6B2-CWZ3E) read by an Arduino Uno run by the Matlab rotary encoder library. Voltage output of the rotary optical encoder was converted to cm, such that 180 (270) cm of virtual track corresponded to 180 (270) cm travel along the treadmill surface. Licks were defined by thresholding voltage from the contact capacitive sensor circuit.

Locomotion traces were first smoothened using a gaussian window of 5 frames. Locomotion stop events were identified as instances when the speed remained below 5 cm/s for longer than 1 second, while locomotion start events were triggered when speeds exceeded 5 cm/s. Events such as CS (conditioned stimuli) were synchronized with the nearest per-plane imaging events occurring at a frequency of 7.8-10.4 Hz. To analyze fluorescent traces around each event, a time window of −5 to 5 seconds was considered. These traces were then averaged to create a single trace per day and normalized by the average value in the period preceding the trigger (−5 to 0 seconds). When calculating CS-triggered averages in both random reward and HRZ (high-reward zone). Early Days were considered to be the first consistent day where the first lick after CS occurred less than 0.25 seconds away and the day prior. Late Days were defined as the last two imaging sessions. These days are combined and averaged for plotting.

Post-CS dopamine amplitudes were determined by averaging values within a 1-second window, with an exception for reward omission trials where the post-CS window was shortened to 400 milliseconds to avoid the period of locomotion difference between rewarded and unrewarded CSs. Ramping was quantified by applying a simple least squares linear fit to the −5 to 0 seconds period of the peri-CS traces. This approach allowed for a comprehensive analysis of dopamine activity and locomotion dynamics surrounding specific events.

## Statistics

Means of data shown with SEM bars or shading. Either one-sample t-tests for difference from one or paired-tests were used. * = p < 0.05; ** = p < 0.01; *** = p < 0.005; **** = p < 0.001; ***** = p < 0.0005.

## Acknowledgements

This work was supported by the National Institute of Mental Health (5R01MH123517), and the McDonnell Institutes for Systems and Cellular/Molecular Neuroscience.

---

**S1.**
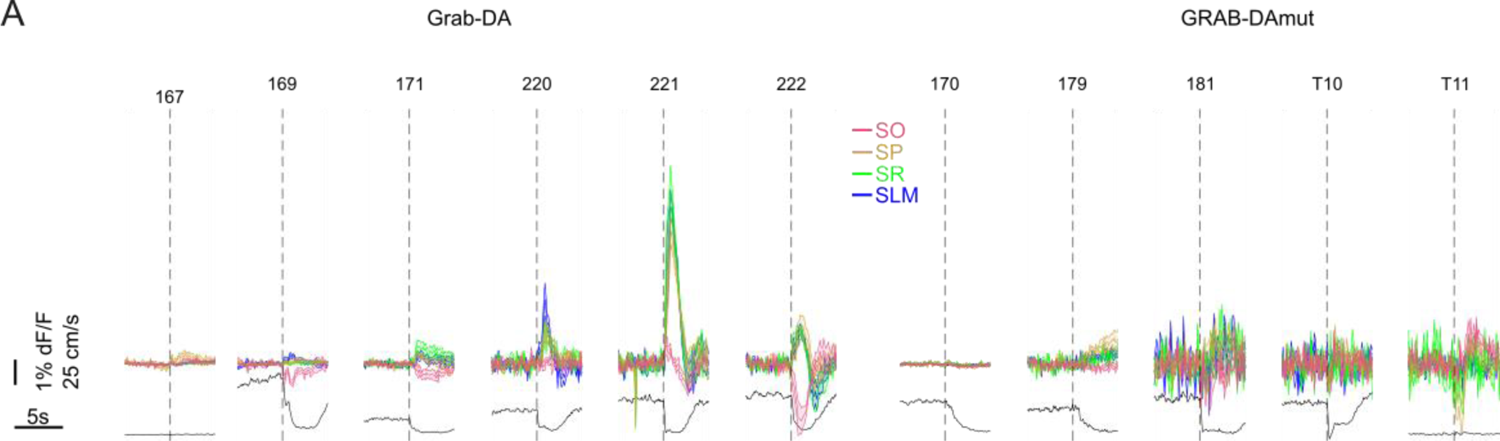
Layer-specific dopamine responses in all other mice. **A**, Early Day CS-triggered average of dopamine fluorescence for all experimental (left) and control (right) mice not shown in Fig 1.

**S2.**
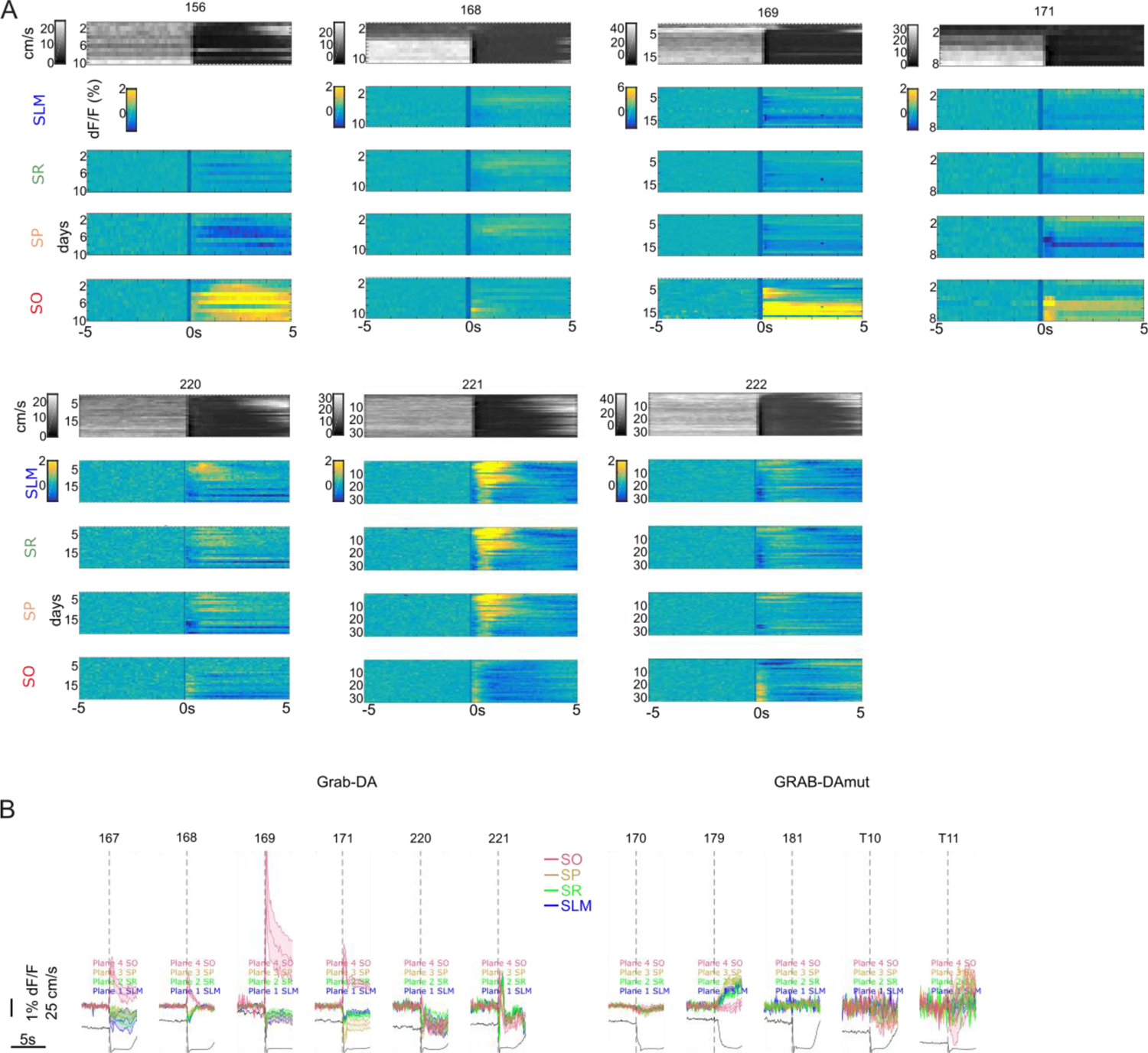
Dopamine relationship with reward and over training. **A,** Speed and dF/F dynamics over days of training in all other experimental mice. **B,** Late Day CS-triggered average of dopamine fluorescence for all experimental (left) and control (right) mice not shown in Fig 2.

**S3.**
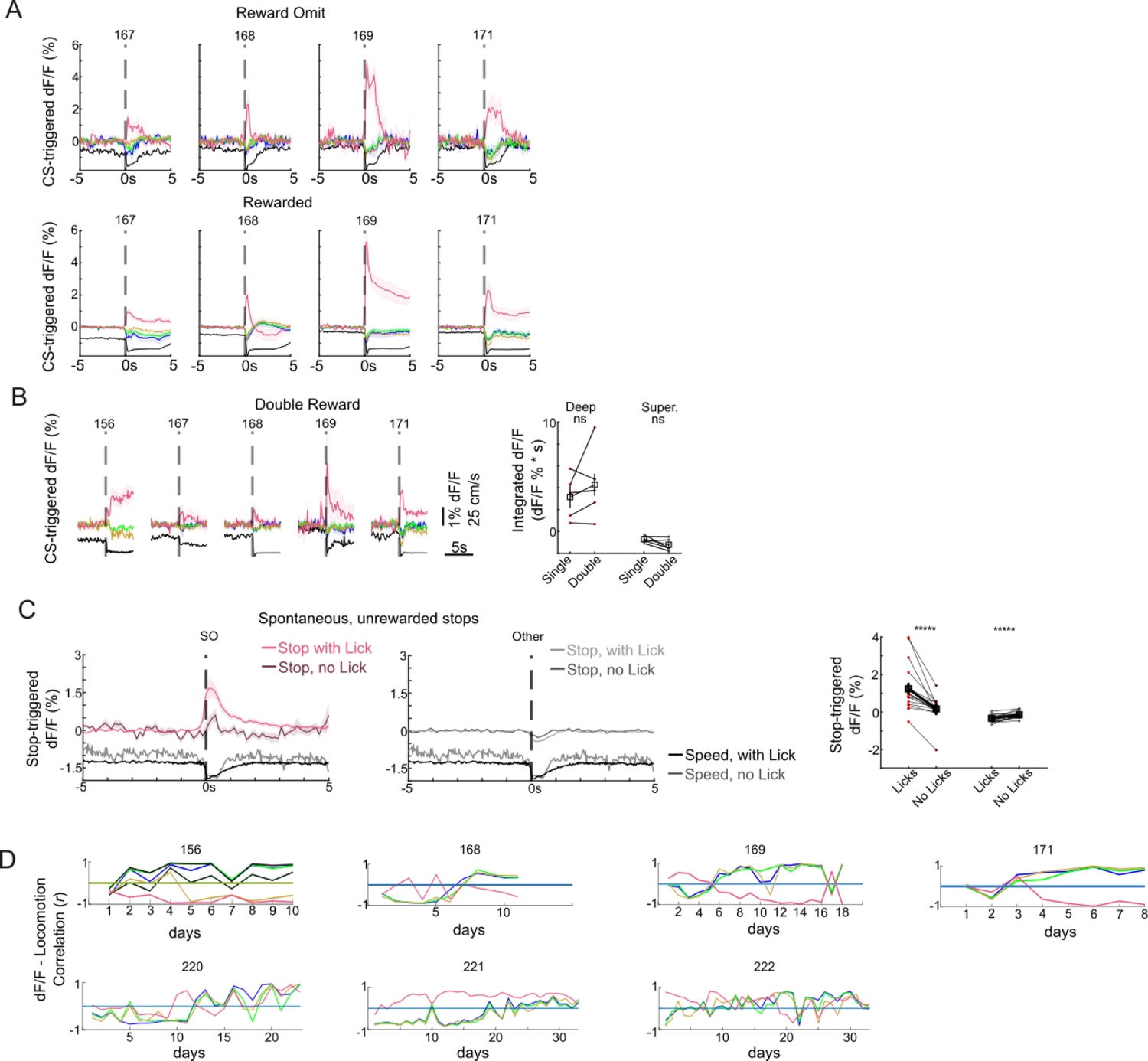
Reward Prediction Error and Locomotion Correlation in all other mice. **A,** CS-triggered average of fluorescence for reward omit (top) and rewarded CS (bottom) from the same imaging sessions for all other GRAB-DA mice. **B,** Left, CS-triggered average of fluorescence for double reward in all GRAB-DA mice. Right, no difference in integrated intensity of CS-triggered dopamine between single and double rewards. **C,** Left, stop-triggered averages, with or without licks (spontaneous, unrewarded stops only). Right, Smaller dopamine transients at stops without licks, n = 14 sessions from 4 mice. **D,** Correlation between plane dF/F and locomotion across training.

**S4.**
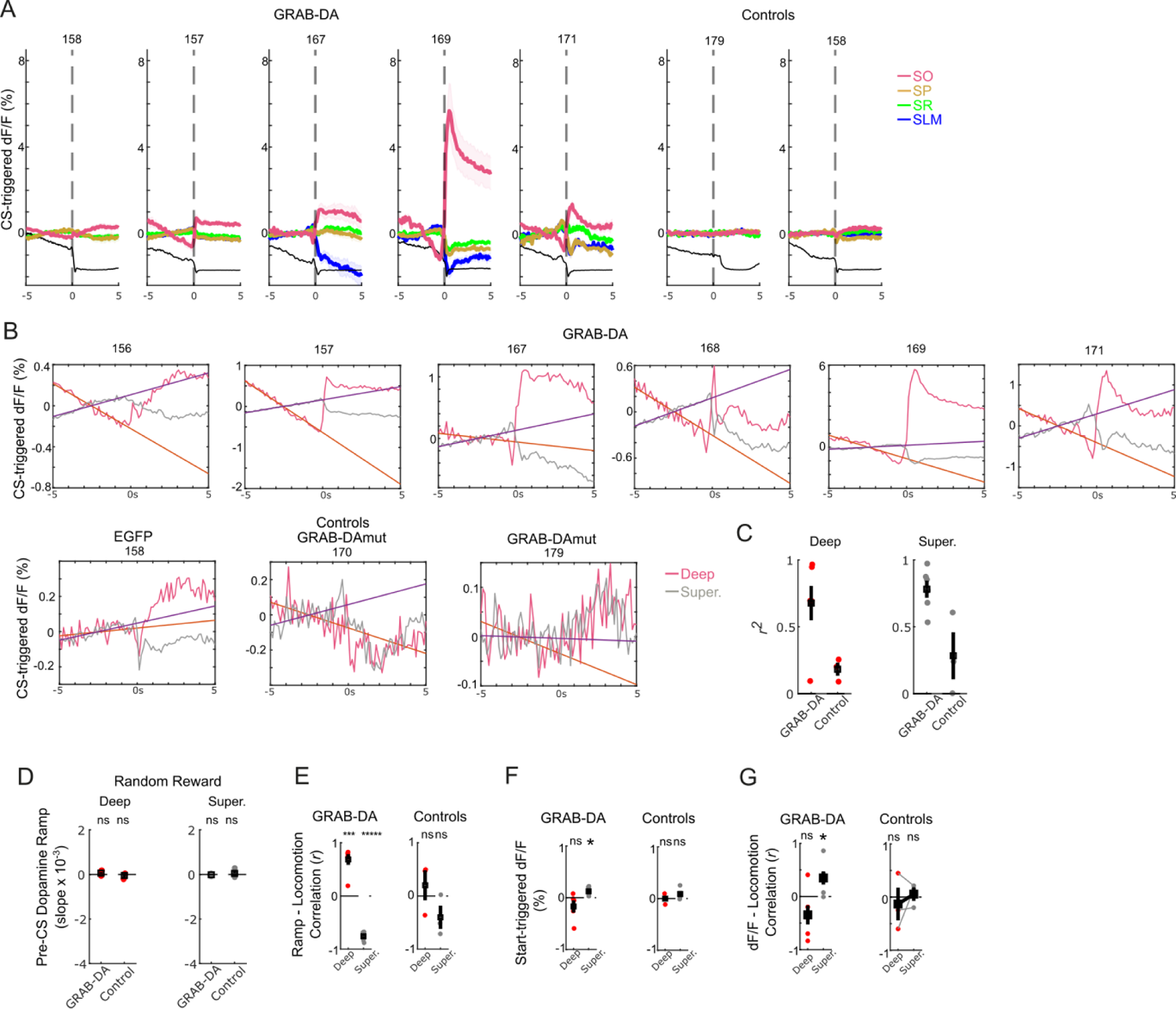
HRZ dopamine transients and ramping. **A,** CS-triggered average from all other mice. **B,** Linear regression of pre-CS dopamine ramps from all mice. **C,** Model fits of linear regression. **D,** No ramping in pre-CS period of random reward behavior. **E,** pre-CS dopamine ramp correlation with locomotion shows correlation of Deep dopamine and anticorrelation of Superficial layer dopaminE, with no significant correlation in controls. **F**, Dopamine in Superficial layers increases with locomotion start. **G**, Dopamine in Superficial layers is significantly correlated with locomotion when the entire time series is considered.

## Supplemental movie

Pseudo-colored movie of dopamine fluorescence intensity in peri-CS window (−5 to +5s) with red showing increased intensity and blue decreased. Scale bar = 18μm.

## Notes

### Competing Interest Statement

The authors have declared no competing interest.

